# CardioSafe: Multi-task prediction of cardiac ion channel activity with reverse-leak audited benchmarking

**DOI:** 10.64898/2026.05.06.723181

**Authors:** M. Jovanović, L. Weidener, M. Brkić, E. Ulgac, A. Meduri

**Affiliations:** Applied Scientific Intelligence, Inc

**Keywords:** cardiotoxicity, hERG, CiPA, multi-task learning, ion channel prediction, cardiac safety, L1000, ChEMBL, molecular fingerprints, Tanimoto similarity

## Abstract

Drug-induced inhibition of the hERG potassium channel is the leading cause of cardiac safety-related drug attrition, but the Comprehensive in Vitro Proarrhythmia Assay (CiPA) framework requires activity data on multiple cardiac ion channels to assess proarrhythmic risk. We present CardioSafe, a three-branch multi-task neural network with cross-attention fusion that integrates chemical fingerprints, ChemBERTa embeddings, and predicted L1000 transcriptomic features to predict blocker status and potency for hERG, Nav1.5, and Cav1.2, with an exploratory IKs head. CardioSafe was trained on the largest publicly reported multi-channel cardiac ion channel dataset, combining ChEMBL 36 with the hERGCentral database (331127 hERG, 3160 Nav1.5, 1138 Cav1.2, and 115 IKs compounds), curated under a pharmacology-aware policy that retains censored measurements and inhibition-percentage votes. Under Tanimoto-similarity-controlled splits, CardioSafe outperforms the leading published comparators (CToxPred2 and CardioGenAI) on the data-rich hERG head; on the smaller Nav1.5 and Cav1.2 heads the standard evaluation is statistically inconclusive. A reverse-leak audit revealed that 22% of Nav1.5 and 21% of Cav1.2 test compounds were present in published comparators’ training data (92% as exact compound matches); after removing these contaminated compounds, CardioSafe’s lead on Nav1.5 and Cav1.2 also reaches statistical significance, demonstrating that prior cross-publication benchmarks for these channels were inflated by training-data overlap.

**Scientific contribution:** We present the first multi-task neural network jointly predicting blocker activity for the three primary CiPA cardiac ion channels (hERG, Nav1.5, Cav1.2) within a single architecture. We introduce a reverse-leak audit methodology that reveals systematic test-set contamination in cross-publication cardiac safety benchmarks, establishing a stricter evaluation protocol. We provide the empirical test of predicted L1000 transcriptomic features as auxiliary input for cardiac ion channel prediction and document a well-characterized negative result.

**Graphical abstract:** CardioSafe encodes each query SMILES with three branches (chemical fingerprints + descriptors, pretrained ChemBERTa, and predicted L1000 transcriptomic signatures), fuses them via a cross-attention block with four learnable per-channel query tokens, and emits binary blocker calls plus pChEMBL regression for hERG, Nav1.5, Cav1.2, and (exploratory) IKs.

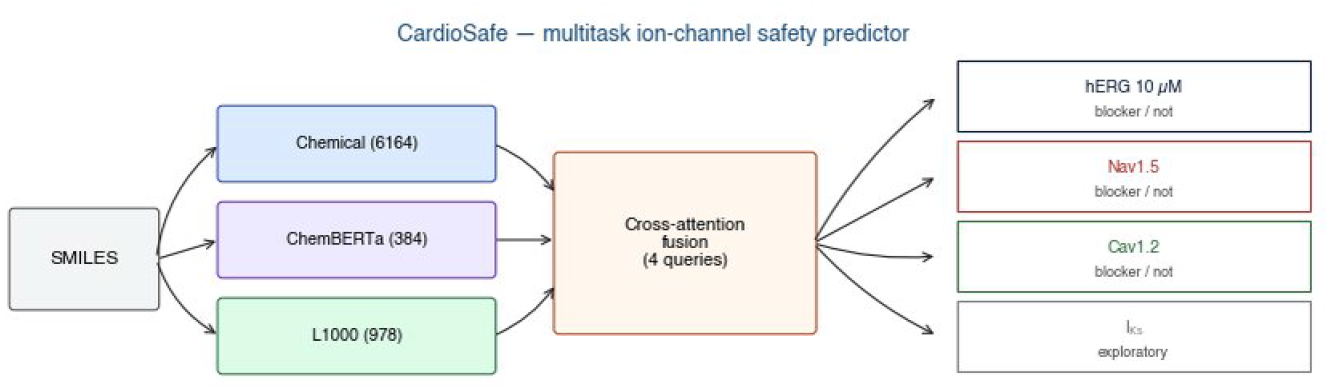

## 1 Background

The human ether-à-go-go-related gene (hERG, *KCNH2*) encodes the Kv11.1 potassium channel, the pore-forming subunit of the rapid component of the delayed rectifier potassium current (IKr) and a major determinant of cardiac repolarization [1]. Drug-induced inhibition of Kv11.1 disrupts repolarization, prolonging the QT interval and elevating the risk of Torsade de Pointes (TdP), a potentially fatal ventricular arrhythmia [1]. Since the first reports linking non-cardiac drugs to TdP in 1965, structurally diverse medications have been shown to induce QT prolongation, leading to market withdrawals such as terfenadine in 1998 [1]. As many as 60% of new molecular entities test positive for IKr blocking liability and are deprioritized in early development [2]. The regulatory framework codified in the International Council for Harmonisation (ICH) guidelines S7B and E14 is narrowly focused on hERG current block and QTc prolongation, both surrogates for proarrhythmia [2]. Although this approach achieves high sensitivity, it has low specificity, causing drugs with little actual TdP risk to be deprioritized [3]. Critically, QT prolongation alone is not sufficient to cause TdP [1]: verapamil is a potent hERG blocker (IC50 = 143 nM) that rarely causes TdP because it concurrently blocks the L-type calcium channel (Cav1.2) at similar potency [1, 2, 4], and amiodarone prolongs QT markedly but carries almost no proarrhythmic propensity owing to its multichannel pharmacology [1]. These examples demonstrate that the net balance of ionic currents, rather than hERG block alone, governs proarrhythmic risk.

The Comprehensive in Vitro Proarrhythmia Assay (CiPA) initiative, proposed in 2013 by the Cardiac Safety Research Consortium, the Health and Environmental Sciences Institute, and the U.S. Food and Drug Administration, formalizes this insight into a regulatory paradigm that evaluates drug effects on multiple cardiac ion channels and integrates them through an in silico cardiomyocyte model [2, 5]. Seven channels were selected for the CiPA panel: IKr (hERG), ICaL (Cav1.2), INa peak and late (Nav1.5), Ito (Kv4.3), IKs (KCNQ1/KCNE1), and IK1 (Kir2.1) [3]. The CiPA in silico model, built on the O’Hara–Rudy human ventricular cardiomyocyte model with a dynamic hERG drug-binding component [6, 7], was prospectively validated on 16 independent drugs with ROC AUC values of 0.89 (manual patch clamp) and 0.98 (hybrid data) [8]. Earlier precursor work had already demonstrated the value of multichannel data: Mirams et al. [9] showed that simulating drug effects on hERG, INa, and ICaL improved TdP prediction above hERG-only markers, and Kramer et al. [10] reported that logistic regression on hERG, Nav1.5, and Cav1.2 IC50 values increased the AUC from 0.77 to 0.93 with a substantial reduction in false positives and false negatives. CiPA has since progressed toward regulatory implementation, with general validation principles codified for proarrhythmia risk prediction models [11, 12, 13]. These developments establish that IC50 data for multiple cardiac ion channels are required inputs to validated regulatory models and motivate machine-learning methods capable of predicting these values at scale.

The central role of hERG in cardiac safety has driven extensive machine-learning model development. Approaches range from attention-based fingerprint classifiers [14] and deep-learning ensembles such as DeepHIT [15] and CardioTox net [16] to graph-based Bayesian models with transfer learning and uncertainty estimation [17], fingerprint-based QSAR with regression capability [18], and graph neural network architectures with dual-level attention [19]. Structure-based approaches have also emerged following the determination of the hERG cryo-EM structure at 3.8 Å resolution [20], with docking-based classifiers achieving AUC values comparable to ligand-based models [21, 22]. Ligand-based approaches often fail in correctly predicting new chemical scaffolds [22], and nearly all published models address hERG alone. Predictive modelling for the remaining CiPA channels is comparatively underexplored. Arab et al. introduced CToxPred [23], providing per-channel classifiers for hERG, Nav1.5, and Cav1.2; their CToxPred2 framework [24] extends this with semi-supervised learning over a large unlabeled chemical space. Kyro et al. [25] subsequently presented CardioGenAI, incorporating discriminative models for the same three channels atop the CToxPred training set. Both train independent per-channel classifiers; no published model has jointly predicted the three primary CiPA channels within a single multi-task framework, nor has any model incorporated biological descriptors beyond chemical structure to compensate for the scarcity of non-hERG training data. IKs prediction has not been attempted from public data at all.

Rigorous evaluation is equally important. Standard benchmarks such as MoleculeNet [26] and the Open Graph Benchmark [27] have shown that scaffold-based splits pose significant challenges for out-of-distribution generalization, and the directed message passing neural network (D-MPNN) architecture, despite strong performance, has yet to reach experimental reproducibility [28]. Activity cliffs cause all methods to struggle, with descriptor-based machine learning outperforming deeper architectures on these compounds [29], and potency prediction benchmarks have been shown to have severe general limitations that prevent reliable method comparison [30, 31]. These findings underscore the need for similarity-controlled evaluation, a concern that is particularly acute for small Nav1.5, Cav1.2, and IKs datasets where a single leaked chemical series can dominate a test set.

One strategy for overcoming data scarcity is to augment chemical representations with biological descriptors derived from gene expression. The L1000 platform measures 978 landmark genes, including the cardiac ion channel genes *KCNH2, SCN5A*, and *CACNA1C*, across over one million profiles spanning roughly 20000 compounds [32]. Integration of gene expression with chemical structure improves adverse drug reaction prediction [33], and transcriptional profiles can identify hERG inhibitors among structurally dissimilar compounds that share toxicity signatures [34]. Deeplearning methods now predict L1000 signatures directly from chemical structure with accuracy that can exceed noisy experimental measurements for downstream tasks [35, 36, 37], and L1000-based models with Bayesian uncertainty have been applied to toxicity prediction [38]. Complementary toxicogenomics resources further support the premise that gene expression data can detect toxicities not observable by conventional assessments [39, 40]. These advances motivate testing whether predicted transcriptomic features can serve as an auxiliary input to cardiac ion channel models, particularly for data-scarce channels, though no such test has been published. Reliable uncertainty estimates are also essential for prospective use: conformal prediction provides a mathematically proven framework with guarantees on error rates and intrinsic applicability domain handling [41], with and large-scale evaluations across hundreds of targets have demonstrated its practical value [42].

In this work, we present CardioSafe, a cross-attention three-branch multi-task neural network that jointly predicts blocker status and potency for hERG, Nav1.5, Cav1.2, and IKs. The chemical fingerprint branch encodes a 6164-dimensional molecular descriptor, the ChemBERTa branch provides a 384-dimensional learned molecular embedding, and the biological branch encodes a 978-dimensional predicted L1000 gene expression signature. We curated the largest publicly reported multi-channel cardiac ion channel dataset from ChEMBL 36, yielding 331127 hERG, 3160 Nav1.5, 1138 Cav1.2, and 115 IKs compounds. All models were evaluated under Tanimoto-similarity-controlled splits at cutoffs of 0.70 and 0.60, and benchmarked against two publicly available comparators, CToxPred2 [24] and CardioGenAI [25], on identical test compounds with a reverse-leak audit. Our objectives were threefold: (1) to empirically test whether predicted transcriptomic features improve multi-channel cardiac ion channel prediction, particularly for data-scarce channels; (2) to establish the first joint hERG / Nav1.5 / Cav1.2 multi-task predictor evaluated under strict similarity-controlled splits, with an exploratory IKs head; and (3) to provide rigorous, de-leaked head-to-head comparisons against existing models on a shared test set. The intended deployment use case is prioritization of compounds for experimental IC50 measurement via patch clamp or microelectrode array (MEA), not regulatory proarrhythmic risk calling, which requires the mechanistic CiPA in silico model.

## 2 Methods

### 2.1 Data curation and preprocessing

We curated ion channel inhibition measurements from ChEMBL version 36 (source dump SHA-256 b25820eef0f0481ad7712bdf4bac3b45 f354e3cbacb76be1fdbf4205d6b48fb9) and supplemented the hERG dataset with the hERGCentral database [43]. Four cardiac ion channel targets were queried: hERG (CHEMBL240, encoded by *KCNH2*), Nav1.5 (CHEMBL1980, *SCN5A*), Cav1.2 (CHEMBL1940, *CACNA1C*), and IKs (CHEMBL2221347, *KCNQ1/KCNE1* complex).

Raw activities were filtered to records `data_validity_comment IS NULL` and drawn from three evidence types: (i) exact potency values with `pchembl_value IS NOT NULL` ; (ii) censored potency values with `standard_relation` in >, >=, <, <=, `standard_units = ‘nM’`, and `standard_type` in IC50, Ki, EC50, Kd, converted to pChEMBL-equivalent via pChEMBL = 9 minus log10(value in nM); and (iii) inhibition percentages (`standard_type = ‘Inhibition’, standard_units = ‘%’`) whose test concentration was parsed from the assay description. SMILES strings were standardized with RDKit 2025.09.6 [44]: parse, salt stripping, charge neutralization, sanitization, and canonical-SMILES regeneration. Each compound was keyed by its standard InChI key.

For each compound and channel, the regression label was the median of exact pChEMBL measurements. The classification label was assigned by a majority-vote rule across all evidence types, with ties resolved to the non-blocker class, using pharmacology-aware rules that differ between the two thresholds. At the 10 µM threshold, a measurement with test concentration at or below 3 µM and greater than 50% inhibition voted blocker (a compound that blocks at low concentration must block at 10 µM); a measurement at exactly 10 µM voted blocker if greater than 50% and non-blocker if at or below 50%; a measurement at 30 µM or above and at or below 50% inhibition voted non-blocker. At the 1 µM threshold, only data recorded at exactly 1 µM contributed a vote. Two classification thresholds were used: pChEMBL > 5.0 (IC50 < 10 µM) and pChEMBL > 6.0 (IC50 < 1 µM). Both thresholds were applied to hERG; Nav1.5, Cav1.2, and IKs were labelled at the 10 µM threshold only.

Per-channel split counts give the number of rows in each split bucket with at least one non-NaN classification label for the channel; per-channel n_train + n_val + n_test equals n_total exactly (e.g. for hERG, 238893 + 46114 + 46120 = 331127). The global compound pool across all four channels comprises 334444 unique standardized compounds, partitioned by the global Tanimoto split into 241792 train, 46326 validation, and 46326 test entries; 753 compounds were removed during split construction to enforce the global Tanimoto constraint. The 1 µM hERG blocker count (3364) is a subset of the 10 µM-labelled hERG compounds with a measurement at the more stringent threshold; Nav1.5, Cav1.2, and IKs are reported at the 10 µM threshold only. The IKs val / test asymmetry (20 / 14) is intrinsic to the global tan70 cutoff together with the forced placement of terfenadine and fexofenadine into the validation fold (see *Data splitting strategy*). The dataset sizes are not directly comparable to those reported by Arab et al. [24] or Kyro et al. [25], because our curation retains censored potency values and pharmacology-aware inhibition-percentage labels that those studies excluded.

### 2.2 Molecular featurization

Each compound was represented by three feature blocks consumed through three separate input branches: a 6164-dim chemical fingerprint-plus-descriptor block, a 384-dim pretrained Chem-BERTa embedding, and a 978-dim predicted L1000 gene expression block. The concatenated input tensor has 7526 dimensions.

#### Chemical fingerprint block (6164 dim)

Three binary fingerprints of length 2048 each were generated with the RDKit `rdFingerprintGenerator` API: (i) Morgan fingerprint with radius 2 (equivalent to ECFP4); (ii) AtomPair fingerprint; (iii) TopologicalTorsion fingerprint. These were concatenated with a 20-dimensional physicochemical descriptor block to form a 6164-dim vector. The descriptor block comprises (i) thirteen RDKit and Gasteiger descriptors: molecular weight, Crippen clogP, topological polar surface area, hydrogenbond donor count, hydrogen-bond acceptor count, rotatable bond count, aromatic ring count, heavy atom count, the mean, maximum, minimum, and standard deviation of Gasteiger partial charges, and the maximum positive Gasteiger charge on any nitrogen atom; and (ii) seven ionization-derived descriptors: the most acidic and most basic predicted pKa values, logD at pH 7.4, and the four microstate fractions (cation, anion, zwitterion, neutral) at pH 7.4. The pKa values were predicted by MolGpKa [45], a graph convolutional neural network ensemble with separate acid and base models, with sentinels of 14.0 and 0.0 used when no ionizable site is detected (the latter applies to 19% of the curated pool, which carry no detectable basic site). logD and the microstate fractions were derived in closed form from clogP and the predicted pKa values using the Henderson– Hasselbalch independent-site model with neutral-microstate-only partitioning. Descriptors were z-scored per-column using means and standard deviations fitted on the training split; zero-variance columns were left unscaled. The full descriptor specification, including per-column non-zero fractions and summary statistics on the production cache, is provided in Supplementary Table S0.

#### ChemBERTa embedding (384 dim)

A 384-dim sentence-level embedding was extracted from the pretrained `DeepChem/ChemBERTa-77M-MTR` transformer [46]. Embeddings were computed once per compound and cached; the transformer was not fine-tuned on our data. This representation provides a learned sequence-level view of the molecule complementary to the handcrafted fingerprints.

#### Biological block (978 dim)

A predicted L1000 gene expression signature across 978 landmark genes, produced by the L1000 expression encoder described in the *Model architecture* section. Predicted signatures were z-scored per-gene using training-split statistics.

### 2.3 Model architecture

We trained CardioSafe, a three-branch multitask neural network with cross-attention fusion (`CrossAttnIonChannelPredictor`), with 3957640 parameters. The model accepts a 6164-dim chemical fingerprint block, a 384-dim pretrained ChemBERTa embedding, and a 978-dim predicted L1000 block (total input dimension 7526).

#### Chemical fingerprint branch

A feed-forward encoder: Linear(6164 →512), BatchNorm1d, ReLU, Dropout(0.2); Linear(512→ 256), Batch-Norm1d, ReLU, Dropout(0.2). The 256-dim output of this branch is used in two places: as input to one of the cross-attention key/value tokens, and as a skip connection concatenated onto every per-channel head.

#### ChemBERTa branch

A single feed-forward projection: Linear(384→ 128), LayerNorm, ReLU, Dropout(0.2). Input embeddings come from the frozen `DeepChem/ChemBERTa-77M-MTR` transformer described in the *Molecular featurization* section.

#### Biological branch

A single feed-forward projection with a higher dropout rate that reflects the higher noise level of predicted transcriptomics: Linear(978→ 128), LayerNorm, ReLU, Dropout(0.3).

#### Cross-attention fusion

Four learnable query tokens, one per ion channel target (hERG, Nav1.5, Cav1.2, IKs), attend to three key/value tokens derived from the three branch outputs. The cross-attention module has attention dimension 128, two attention heads, and attention dropout 0.1. Query tokens were initialized as `torch.randn(4, 128) * 0.02.` The attention output for each channel is a 128-dim context vector. The two hERG classification heads (10 µM and 1 µM) and the hERG regression head all consume the same hERG query-token context vector; analogous query-to-head sharing applies for Nav1.5 and Cav1.2 (one classification head and one regression head each), and the IKs query token feeds only the single IKs classification head.

#### Heads

Eight task-specific heads were implemented identically. Each head concatenated the per-channel 128-dim context vector with the 256-dim chemical-branch skip connection (total 384 dim) and passed through Linear(384→128), ReLU, Dropout(0.3), Linear(128→1). Three regression heads predicted pChEMBL for hERG, Nav1.5, and Cav1.2. Five classification heads predicted binary blocker labels for hERG at 10 µM, hERG at 1 µM, Nav1.5, Cav1.2, and IKs. IKs had no regression head given insufficient continuous measurements.

#### Initialization

All Linear layers were initialized with Kaiming-normal weights (ReLU nonlinearity) and zero biases, except the four learnable query tokens described above. Classification heads emitted raw logits; a sigmoid was applied at inference to produce blocker probabilities. Regression targets were z-scored per-channel with training-split statistics and inverse-transformed at evaluation.

#### L1000 expression encoder

The biological branch’s input was generated by a separately trained neural network that mapped a 1024-dimensional TopologicalTorsion count fingerprint to 978 predicted landmark-gene z-scores. The encoder consisted of a gene co-expression graph convolutional network (GCN) with two GCNConv layers on 384-dimensional gene embeddings (edges retained where the absolute Pearson correlation of training-set signatures exceeded 0.4), a shared three-layer feed-forward network of width 512 with ReLU and dropout of 0.2 between the concatenated molecular and gene representations, and per-gene linear heads. The encoder was trained on Level-5 consensus signatures (MODZ) aggregated from LINCS Phase I (GEO accession GSE92742) and Phase II (GEO accession GSE70138), restricted to small-molecule perturbations (`pert_type = trt_cp`) and compounds with at least five replicate signatures. Signatures were averaged across dose, cell line, and timepoint for each compound prior to encoder training. The L1000 co-expression graph threshold (|r|) was tested at 0.30, 0.40, 0.50; downstream CardioSafe AUC varies by less than 0.005 on hERG across all three thresholds. The deployed |r| = 0.40 matches the threshold at which CMap researchers report typical co-expression filtering.

Averaging across cell lines is a deliberate choice forced by the LINCS L1000 panel itself: across Phase I and Phase II, the panel covers 71 and 30 unique cell lines, respectively, none of which is of cardiac origin. The model therefore cannot access cardiac-tissue-specific transcriptomic context, and the biological-branch signal that it does exploit is a cross-tissue perturbation signature. The implications of this tissue-averaging choice are discussed in the *Discussion*.

### 2.4 Training procedure

#### Two-stage training

CardioSafe was trained in two stages. Stage 1: the full model was trained end-to-end with focal loss (gamma = 2.0, per-head positive-class weight capped at 8) using the Adam optimizer with weight decay 1e-5, a Noam-style learning rate schedule (linear warmup from 1e-4 to 1e-3 over 3 epochs, followed by exponential decay to 1e-5), batch size 256, maximum 100 epochs, and early stopping with patience 20 on hERG 10 µM validation AUC.

Stage 2 (cliff fine-tune) continued training each of the 5 seed checkpoints from Stage 1 for 9 epochs of 120 mini-batches each at a constant learning rate of 1e-5, using AdamW with weight decay 1e-5 and gradient clipping at L2 norm 1.0. The cliff training set was assembled in two curation stages. First, a manual literature curation across 25 published cardiac-cliff sources (anchored on Bowes et al. [47], Kramer et al. [10], and 23 smaller-source contributions detailed in Supplementary Note S2) yielded 53 compounds organized into 30 therapeutic-class `pair_id` groups in which clinical hERG-safety divergence is established between a blocker member and a safer analogue (categories include antiarrhythmic, antihistamine, atypical antipsychotic, beta-blocker, butyrophenone, fluoroquinolone, gastroprokinetic, opioid, phenothiazine, and tricyclic antidepressant). Selection was based on documented clinical-safety divergence within a therapeutic class; no automated Tanimoto-similarity or Δ-pIC50 threshold was applied. Second, an automated leak-prevention filter dropped any cliff compound whose Morgan radius-2 2048-bit Tanimoto to any validation or test row of the active split exceeded the split’s cutoff (0.70 on tan70; 0.60 on tan60), guaranteeing that no information from training cliff compounds leaked into the held-out evaluation set. After filtering, 48 cliff compounds across 29 `pair_id` groups remained on tan70 (terfenadine, fexofenadine, ranolazine, desipramine, and amitriptyline were dropped as they sit in the validation fold), and 51 compounds across 29 `pair_id` groups remained on tan60 (only terfenadine and fexofenadine were dropped, as the other three sit in the train fold under the 0.60 cutoff). Of the surviving `pair_id` groups, 12 (tan70) or 13 (tan60) contain both a blocker and a safer member and contribute (blocker, safer) ranking pairs to the loss; the remainder contain only one role.

Each Stage 2 mini-batch concatenates 512 randomly sampled ChEMBL training rows with all retained cliff compounds at an 8-fold sample weight. The optimization target is the same multi-task focal-BCE (gamma = 2.0) plus regression-MSE used in Stage 1, augmented with a pairwise margin-ranking loss `max(0, 1.5 $-$ (logit_blocker $-$ logit_safer))` with coefficient 0.3, applied only to the two hERG classification heads (hERG 10 µM and hERG 1 µM) since the cliff curation is anchored on hERG safety. The ranking term explicitly pushes the blocker member of each cliff pair above its structurally similar safer counterpart in logit space. All eight task heads receive gradient through the multi-task term; the ranking term influences only the two hERG heads. Cross-attention queries, descriptor and L1000 scaler statistics, and ChemBERTa weights are inherited unchanged from the Stage 1 checkpoint. Stage 2 wall-clock time is approximately 5 min for all 5 seeds run in parallel.

**Figure 1:**
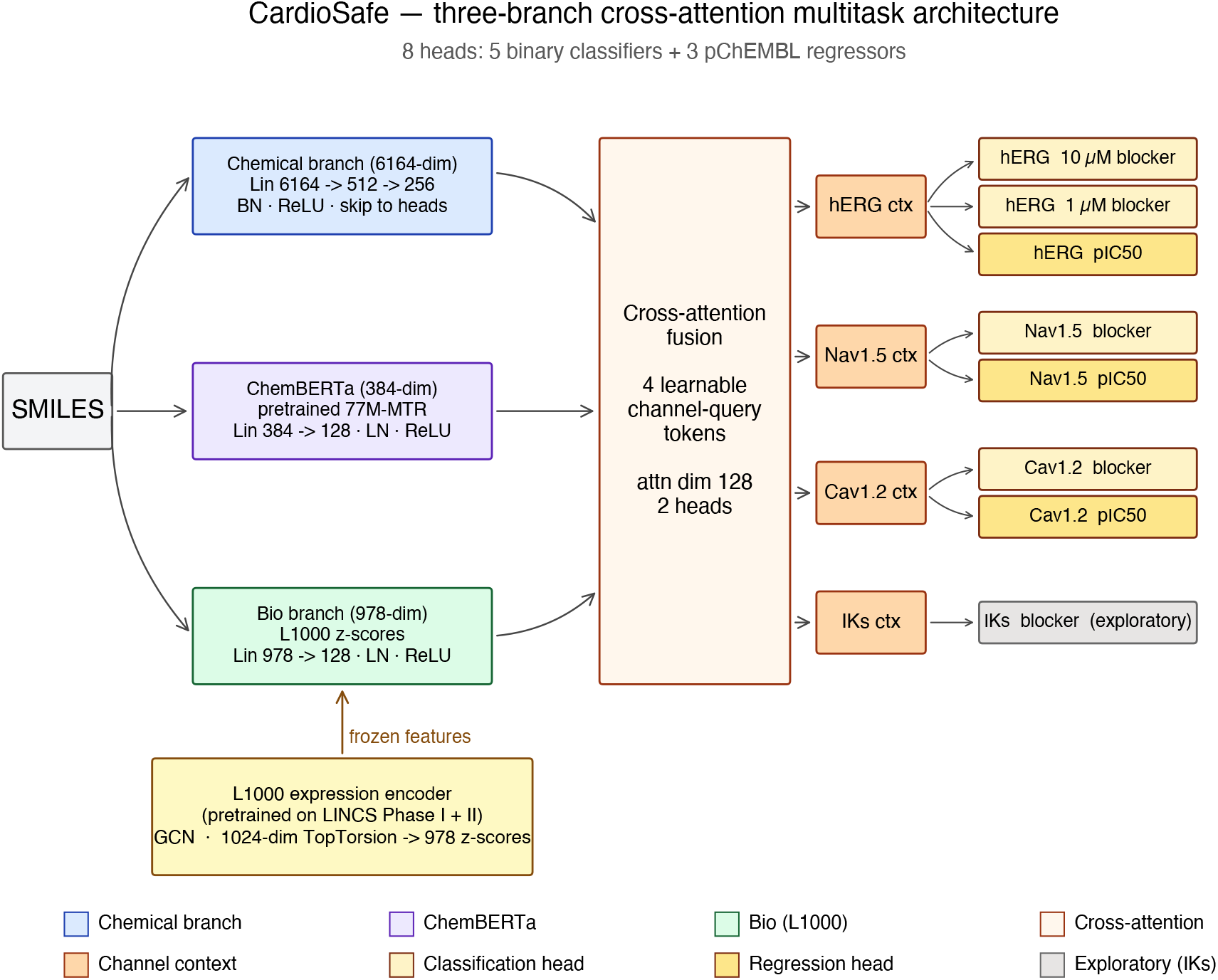
Architecture of CardioSafe. Three input branches encode a 6164-dim chemical fingerprint-plus-descriptor block, a 384-dim pretrained ChemBERTa embedding, and a 978-dim predicted L1000 transcriptomic signature (the L1000 encoder is a separately trained graph convolutional network mapping a 1024-dim TopologicalTorsion count fingerprint to landmark-gene z-scores). A cross-attention fusion layer with four learnable per-channel query tokens (hERG, Nav1.5, Cav1.2, IKs) attends over the three branch outputs to produce per-channel context vectors. Each of the eight task heads (five binary classifiers, three pChEMBL regressors) consumes its per-channel context vector concatenated with a 256-dim skip connection from the chemical branch. The IKs head (shaded) is exploratory.

#### Multi-seed training

Five independent training runs were executed with random seeds 42 through 46. Primary metrics are reported as the 5-seed ensemble (average of sigmoid probabilities for classification, average of inverse-scaled predictions for regression) with bootstrap 95% confidence intervals (1000 resamples).

#### Hardware and runtime

Training was per-formed on an Apple M4 Pro (12-core CPU, 24 GB unified memory) using PyTorch MPS. Per-seed wall-clock time was 36–49 min (median 44 min); the full 5-seed ensemble required approximately 3.7 h.

#### Software

PyTorch 2.10.0 [48], PyTorch Geometric 2.7.0 (L1000 GCN), RDKit 2025.09.6 [44], scikit-learn 1.8.0 [49], Python 3.13.6. ChemBERTa embeddings were computed with the `DeepChem/ ChemBERTa-77M-MTR` checkpoint [46].

#### Statement of generative AI usage

Claude Opus 4.7 (Anthropic) was used to refine the language and grammar. All scientific content, results, and conclusions are the responsibility of the authors.

### 2.5 Data splitting strategy

#### Primary split (tan70)

Our primary evaluation used an in-house Tanimoto-similarity-controlled split at cutoff 0.70, referred to as *tan70* throughout this paper. For each ion channel, every labelled compound’s Morgan fingerprint (radius 2, 2048 bits, as used in the chemical branch) was compared against all other labelled compounds for that same channel, and a graph of pairwise Tanimoto similarities at or above 0.70 was assembled. Connected components of this graph were partitioned into train, validation, and test buckets with an 80/10/10 target ratio, under the constraint that no cross-bucket edge with similarity at or above 0.70 is allowed. Partitioning was verified post-hoc by recomputing the full cross-bucket similarity matrix; zero violations at or above the 0.70 cutoff were observed. The resulting test pool contains 46326 compounds across all heads. Both splits force terfenadine and fexofenadine into the validation fold to keep them out of any seed’s training pool, enabling their use as a held-out case study (see *Results*).

#### Secondary split (tan60)

A second, stricter split was constructed identically at Tanimoto cutoff 0.60 and is reported where it provides additional information (for example, performance degradation under stricter novelty). The tan60 test pool is smaller (13889 compounds) and its per-channel positive counts are limited (IKs: 1 blocker out of 11 test compounds).

#### Why not Bemis–Murcko scaffold split

Bemis– Murcko scaffold splitting is conceptually attractive but penalizes chemical-series diversity unevenly across channels. Tanimoto-similarity splits at a fixed cutoff directly guarantee that no test-set compound shares a near-neighbour, as defined by the same fingerprint used by the model, with any training compound, which is the property we care about for external-generalization claims.

### 2.6 Evaluation metrics

Classification and regression heads were evaluated independently for each ion channel, with at least 5 test-set compounds required for a head to be scored. Metrics were computed on the unscaled original pChEMBL scale: regression predictions were inverse-transformed using the training-split scaler before any metric was computed.

#### Classification metrics

For every binary head we computed: area under the receiver operating characteristic curve (AUC-ROC), undefined and omitted when only one class was present in the test split; Matthews correlation coefficient (MCC); sensitivity; specificity; accuracy; and area under the precision–recall curve (AUPRC). MCC is preferred over F1 as the primary summary statistic because it is symmetric in the two classes and remains informative at the 2% blocker prevalence of our largest evaluation set; F1 is dominated by rare-positive recall at this prevalence and is not reported.

#### Regression metrics

For each regression head we computed root mean square error (RMSE), mean absolute error (MAE), Pearson correlation coefficient (*r*), and Spearman rank correlation (rho). We report Pearson *r* as the primary regression metric and Spearman rho alongside because the latter is invariant to the prediction-scale compression observed on the stricter tan60 holdout.

#### Uncertainty quantification

All primary metrics are reported as the 5-seed ensemble point estimate with bootstrap 95% confidence interval (1000 resamples of test-set triples with replacement, reporting the 2.5th and 97.5th percentiles). For head-to-head comparisons against comparator models, a paired bootstrap was used: the same bootstrap indices were applied to both sets of predictions, yielding a 95% CI for the difference and a one-sided exceedance probability. The 5-seed mean *±* standard deviation is reported alongside for seed-sensitivity transparency.

#### External validation and comparator analysis

We positioned CardioSafe against CToxPred2 [24] and CardioGenAI [25], both of which accept SMILES input and predict hERG, Nav1.5, and Cav1.2 classification outputs at matched thresholds. CardioGenAI uses the same training set as CToxPred (per-channel, verbatim), so the leakage audit applies identically to both comparators. For each comparator, we generated predictions on the full tan70 test pool using the comparator’s released code and default checkpoints. Reported numbers therefore use identical test compounds, identical ground-truth labels, and identical metric definitions across all models.

### 2.7 Y-randomization control

One full training run was performed on each split (seed 42) after permuting all classification and regression labels within the training set. All classification MCC values collapsed to within *±*0.07 of zero (hERG 10 µM: 0.043; hERG 1 µM: *™*0.022; Nav1.5: 0.050; Cav1.2: *™*0.062; IKs: 0.000), and all regression Pearson *r* values collapsed (hERG: 0.104; Nav1.5: 0.019; Cav1.2: *™*0.288).

The hERG 10 µM Y-randomized AUC was elevated above 0.5 (0.70 on tan70; 0.68 on tan60) due to a residual correlation between molecular weight and the original blocker label that survives per-column permutation under the natural 2% positive prevalence; we provide a stratified analysis in Supplementary Note S1 confirming the AUC collapses to 0.50–0.62 within molecular-weight strata. The regression-*r* and MCC collapses (the unconfounded controls) confirm no information leakage.

## 3 Results

All primary numbers are the 5-seed ensemble point estimate with bootstrap 95% confidence interval (1000 resamples). Both tan70 (primary) and tan60 (secondary, stricter) splits are reported.

### 3.1 Classification performance

The hERG 10 µM head achieved stable generalization to structurally novel compounds, with tight inter-seed variation (5-seed standard deviation 0.002 on tan70). The hERG 1 µM head, aligned with the CiPA clinical regime, achieved higher AUC (0.959), consistent with more potent blockers being easier to separate from non-blockers. Nav1.5 and Cav1.2 achieved AUC values of 0.831 and 0.841 on tan70 despite substantially smaller training pools.

For IKs, where only 115 compounds with 30 blockers were available, we report a first publicly evaluated prediction head (n = 14 test compounds, 1 blocker on tan70; n = 11, 1 blocker on tan60). Performance is statistically indistinguishable from random: the bootstrap 95% CI on AUC includes 0.5 in both splits, and we present this result to motivate further IKs data curation rather than as a performance claim.

At the 2.05% blocker prevalence of the hERG 10 µM test pool, an MCC of 0.47 indicates that the model produces false positives at the default 0.5 threshold. In a safety screening context this is an acceptable trade-off: the model is most useful for prioritizing compounds for experimental IC50 measurement, and threshold adjustment can shift the sensitivity/specificity balance to match the cost asymmetry of a given screening campaign.

**Figure 2:**
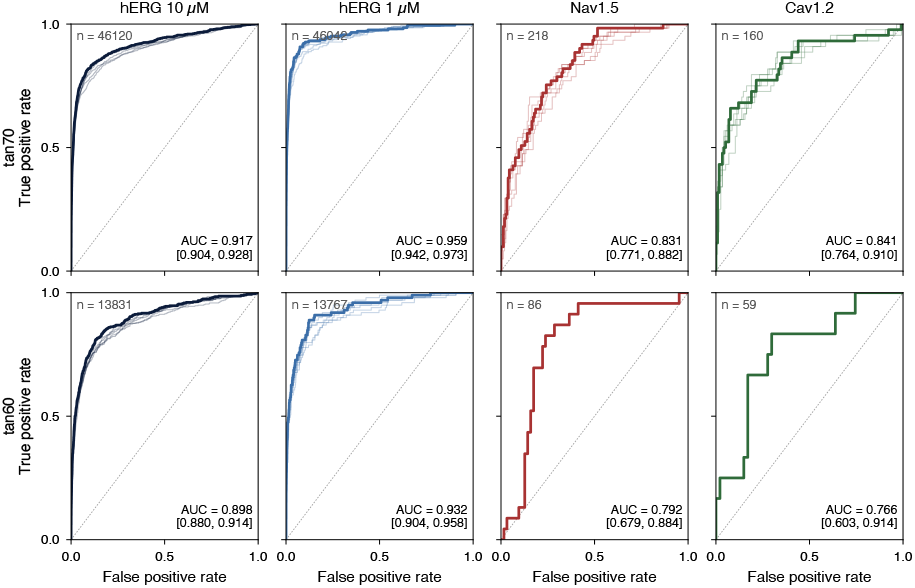
Receiver operating characteristic curves for CardioSafe across the four cardiac ion channel classification heads, evaluated on the primary tan70 split (top row) and the stricter tan60 split (bottom row). Bold curves show the 5-seed ensemble; faint overlaid curves show individual seeds (n = 5). Annotations report ensemble AUC with bootstrap 95% confidence intervals (B = 1000 resamples, seed 20260428). The IKs head (n = 14 / 11 test compounds, 1 blocker) is omitted as bootstrap CIs include AUC = 0.5 on both splits.

### 3.2 Regression performance

Pearson *r* drops by 0.12–0.36 from tan70 to tan60 on every head, as expected from the stricter Tanimoto cutoff. Spearman rho holds up better than Pearson on Nav1.5 tan60 (0.580 vs. 0.197), indicating the rank ordering is preserved but the prediction scale is compressed. On tan60 hERG and Nav1.5, *R*^2^ turns negative (predictions worse than predicting the test mean in absolute terms), consistent with ensemble predictions compressing around the training mean while the test distribution has heavier tails.

### 3.3 Comparator benchmarking

We compared CardioSafe directly against CToxPred2 [24] and CardioGenAI [25] on identical compounds: predictions from each comparator’s released code and default checkpoints were generated on the same tan70 test pool. CardioGenAI uses the same training set as CToxPred (per-channel, verbatim), so the leakage audit applies identically to both comparators. Paired-bootstrap differences (B = 1000 resamples, common indices applied to both models, random seed 20260428) were computed for every (channel, comparator) pair on both metrics.

#### Reverse-leak audit

Because all models draw from overlapping ChEMBL records, a fraction of our test pool was inevitably present in comparator training data. This is a structural property of the subfield rather than a procedural error by any individual study. Two thresholds were used in the audit. The broad audit at Tanimoto ≥ 0.70 identifies any near-neighbour overlap: 3.1% of hERG test compounds (1424 of 46120), 22.0% of Nav1.5 test compounds (48 of 218), and 20.6% of Cav1.2 test compounds (33 of 160) had a nearest comparator-training neighbour at this threshold. Of these near-neighbours, 92% were exact InChI-key matches; only 0.3% of hERG test compounds had fuzzy near-neighbours (Tanimoto 0.70–0.99) in the comparator training pool. The leak is therefore dominated by exact compound overlap rather than scaffold spillover. For the de-leaked head-to-head comparison, we used the stricter Tanimoto ≥ 0.99 threshold, which removes the dominant exactmatch contamination component (2.8% of hERG test, 22.0% of Nav1.5 test, and 20.6% of Cav1.2 test compounds) while retaining the 0.3% of hERG test compounds that share scaffolds but are not identical molecules.

#### Head-to-head comparison

Tables 3 and 3b summarise the pre-de-leak (full per-channel valid test fold) and post-de-leak (test fold minus comparator-train compounds at Tanimoto ≥ 0.99) head-to-head performance.

Three reads of these tables are worth noting. First, on the data-rich hERG 10 µM head (n ≈ 46k), CardioSafe wins both AUC and MCC against both comparators by margins whose 95% CIs exclude zero by a wide margin in both pre-and post-de-leak views. The post-de-leak Δ-AUC widens (from +0.085 to +0.131 vs CToxPred2; from +0.055 to +0.115 vs CardioGenAI), consistent with CardioSafe’s hERG ranking having been correct on the leaked compounds while the comparators recovered them partly through training memorization. Second, on the smaller Nav1.5 (n = 218) and Cav1.2 (n = 160) heads the pre-de-leak Δ-AUC CIs straddle zero, but the post-de-leak Cav1.2 Δ-AUC vs CToxPred2 (+0.106 [+0.032, +0.194]) and Nav1.5 Δ-AUC vs CardioGenAI (+0.075 [+0.003, +0.146]) clear zero. Third, the most informative single observation is the pre-versus-post-de-leak shift on Cav1.2: CardioGenAI is ahead by 0.032 AUC pre-de-leak and behind by 0.022 AUC post-de-leak. This flip, driven by the removal of 33 of 160 test compounds that appear verbatim in CardioGenAI’s training set, is the cleanest illustration that prior cross-publication Cav1.2 head-to-head numbers in this subfield were inflated by training-data overlap.

CardioSafe’s primary advantages over comparators are therefore on Nav1.5 and Cav1.2 (where de-leaking reveals the largest corrections) and on the evaluation methodology itself, rather than on hERG, where the model wins decisively but other strong published baselines exist.

This issue is structural rather than procedural: all cardiac ion channel models draw from overlapping ChEMBL records, and no established protocol exists for auditing cross-publication compound overlap. Our reverse-leak analysis is intended to fill that methodological gap rather than to criticize the affected models, which followed the evaluation norms standard at the time of their publication. We recommend that both InChI-key intersection audits and Tanimoto nearest-neighbour audits become standard practice in cardiac ion channel benchmarking.

### 3.4 Branch ablation

We quantified the contribution of each auxiliary branch by training CardioSafe with (a) the L1000 biological branch held to zero (bio-zero) and (b) the ChemBERTa embedding held to zero (chemberta-zero), using the same 5 seeds, splits, and training schedule.

The L1000 biological branch contributes no measurable improvement to headline classification performance. AUC deltas between deployed and bio-zero sit within *±*0.008 on tan70 for all heads with n ≥ 86; the 95% confidence intervals overlap heavily in every case. On the stricter tan60 split, removing the bio branch nominally improves Cav1.2 binary classification (deployed AUC 0.766 vs. bio-zero 0.801, Δ +0.035), although the CIs again overlap.

The ChemBERTa branch shows a more nuanced pattern. On tan70 hERG 10 µM, removing Chem-BERTa drops AUC from 0.917 to 0.904, the largest deployed-versus-ablation gap on any classification head, although the 95% CIs overlap modestly (deployed [0.905, 0.929] vs. ChemBERTa-zero [0.892, 0.916]). The deployed model’s lower bound exceeds the ablation point estimate, suggesting a small but consistent benefit. On every other head the CIs overlap substantially.

We retain both auxiliary branches in the deployed model because (a) no single ablation wins on every head, (b) the parameter and latency cost is dominated by the chemical branch, and (c) future integration of cardiac-specific transcriptomic data could change the bio-branch finding. However, neither branch is load-bearing for headline AUC, and the chemical fingerprint branch carries the model’s discriminative signal.

#### Cross-attention introspection

Analysis of the learned query tokens and attention weights revealed that the cross-attention layer operates at near-uniform entropy across all four channel queries (1.098 nats versus the uniform-3-key maximum of log(3) = 1.099 nats). No channel has learned to specialize its attention toward a preferred branch. The per-channel head outputs are dominated by the 256-dim chemical-branch skip connection, not the 128-dim attention context. The cross-attention specialization that the architecture was designed to learn does not materialize at this data scale; the model behaves as a single-trunk multi-task chemical MLP with a small uniform-attention bonus.

### 3.5 Curation-policy sensitivity

Restricting training to strictly exact pIC50 measurements (`standard_relation = ‘=’`, matching the criterion of Arab et al. [23]) reduces hERG binary AUC by 0.18 on tan70 (0.917 to 0.737) and 0.21 on tan60 (0.898 to 0.688). hERG drops 93% of binary labels under this restriction because the majority of hERG binary labels in ChEMBL derive from inhibition-percentage or censored-measurement votes, not exact IC50. Nav1.5 and Cav1.2 binary AUC drops are smaller (0.04–0.07) because those channels lose only 10% of training rows.

Regression accuracy on tan60 improves under the strict exact-only policy (hERG *r*: 0.331 to 0.443; Nav1.5 *r*: 0.197 to 0.325), confirming that the noisy non-exact measurements help classification but introduce per-compound noise that hurts regression on harder out-of-distribution splits.

### 3.6 Applicability domain analysis

For every tan70 test compound, we computed Tanimoto similarity to its nearest training-set neighbour and binned the test pool into three strata.

**Table S7 (excerpt)** Per-bin classification performance on tan70

hERG 10 µM AUC degrades from 0.920 in the 0.5–0.7 bin to 0.812 in the 0.0–0.3 bin (0.108 drop), and MCC drops from +0.518 to +0.218. Critically, the hERG 1 µM head at the 0.0–0.3 stratum maintains AUC of 0.844 (reasonable rank ordering) but MCC collapses to 0.003 (the operating-point classifier fails). AUC alone underestimates operational risk at low structural similarity to training data. Below Tanimoto *™*0.30 the deployed classifier should not be relied upon at the default threshold.

### 3.7 Failure-mode analysis

The top-20 most-confident false predictions per classification head were categorized into five buckets. Across both splits and all five heads (269 false predictions examined):

Notably, only 3% of the most-confident failures occur outside the applicability domain (max Tanimoto < 0.30); the model is not making confident wrong calls out of domain but rather making low-confidence calls that do not dominate the top-confident-wrong list. Prodrug motifs (12%) are the clearest expected-SAR mismatches: the model rates the prodrug parent as a blocker because active-form analogues are blockers, but the prodrug itself is rapidly hydrolyzed in vivo. Label noise (2%) includes physically impossible hERG label inconsistencies (1 µM = blocker but 10 µM = non-blocker). The two largest tractable buckets, prodrug motifs and label noise, admit straightforward mitigations: prodrug-aware standardization that resolves to the active metabolite before featurization, and a label-consistency filter that flags the small subset of compounds with cross-threshold contradictions for review.

### 3.8 Terfenadine/fexofenadine case study

Both compounds were forced into the validation fold (unseen by all 5 seeds during training). On tan70:

CardioSafe correctly resolves the qualitative direction and clinical call of this canonical hERG activity cliff: terfenadine is correctly classified as a strong blocker (predicted pIC50 6.38, blocker classification output (CO) 0.895 at 10 µM and 0.763 at 1 µM) and fexofenadine as a borderline non-blocker (predicted pIC50 4.80, blocker CO 0.683 at 10 µM and 0.389 at 1 µM). The predicted pIC50 gap of +1.58 underestimates the labelled cliff (terfenadine pChEMBL = 7.55, fexofenadine = 3.96; labelled cliff = +3.59); the model preserves the rank ordering and crosses the operational thresholds correctly, but compresses the activity range, a pattern consistent with the regression-mean-reversion behaviour observed on tan60 (Table 2b). Inter-seed standard deviations are low for terfenadine (0.020 on hERG 10 µM) and moderately higher for fexofenadine (0.029), reflecting the latter’s proximity to the decision boundary.

**Figure 3:**
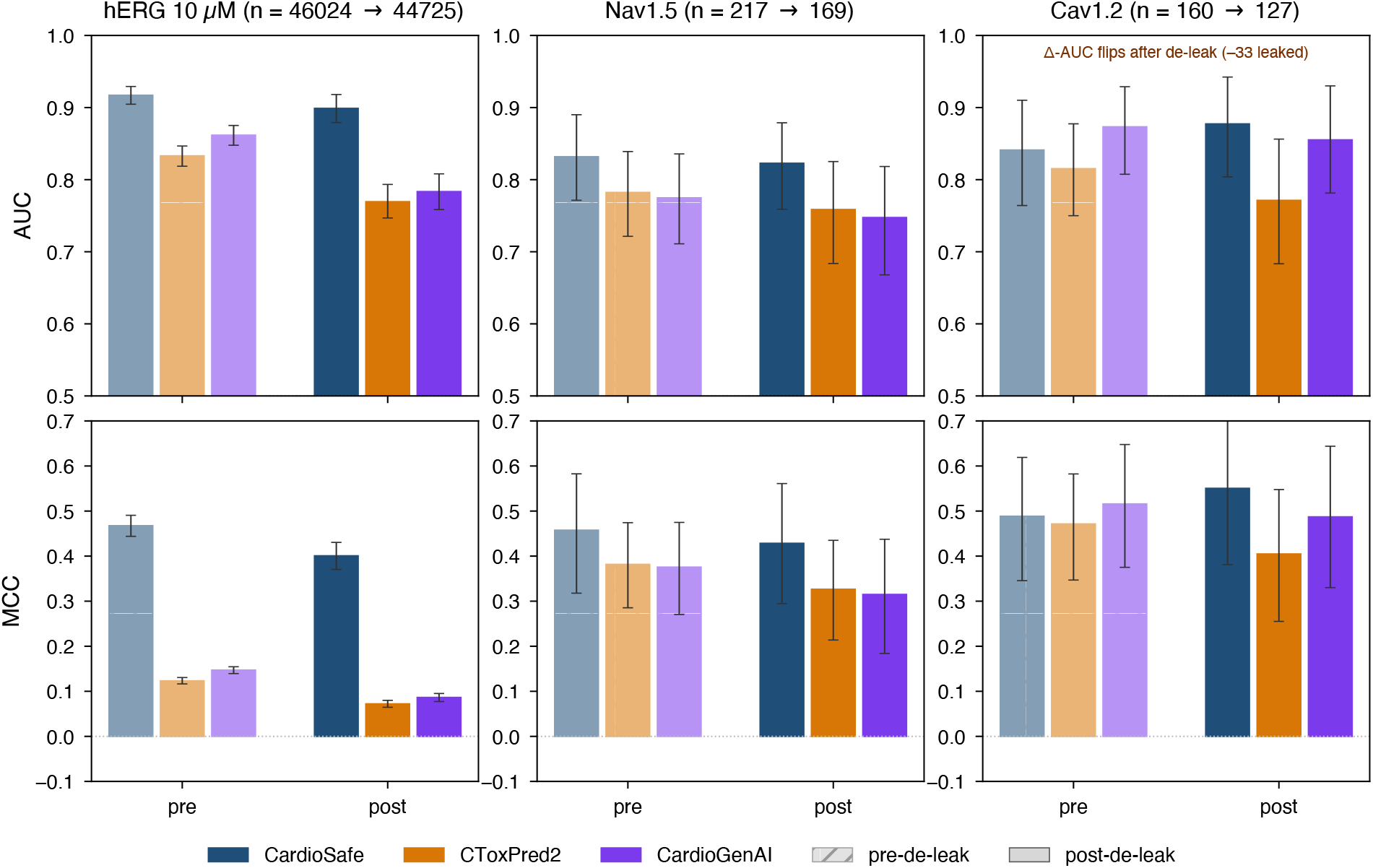
Head-to-head comparison of CardioSafe against CToxPred2 and CardioGenAI on the tan70 test fold, before and after removing compounds present in the comparators’ training data at Tanimoto ≥0.99 (the dominant exact-match component of cross-publication contamination). Top row: AUC; bottom row: Matthews correlation coefficient. Hatched bars are pre-de-leak; solid bars are post-de-leak. Error bars are bootstrap 95% confidence intervals (B = 1000 paired resamples, common indices applied to all models). The Cav1.2 panel illustrates the largest de-leak correction: pre-de-leak CardioGenAI is ahead of CardioSafe by Δ-AUC = *™*0.032; post-de-leak (with 33 of 160 test compounds removed as exact matches to the comparator training pool) CardioSafe is ahead by Δ-AUC = +0.022.

**Table 1:** Dataset composition per channel (post-curation pool, labels_v1)

| Channel | n_total | n_blockers (10 μM) | n_blockers (1 μM) | n_train (tan70) | n_val (tan70) | n_test (tan70) |
| --- | --- | --- | --- | --- | --- | --- |
| hERG | 331127 | 11881 | 3364 | 238893 | 46114 | 46120 |
| Nav1.5 | 3160 | 1240 | n/a | 2689 | 253 | 218 |
| Cav1.2 | 1138 | 548 | n/a | 833 | 145 | 160 |
| IKs | 115 | 30 | n/a | 81 | 20 | 14 |

**Table 2:**
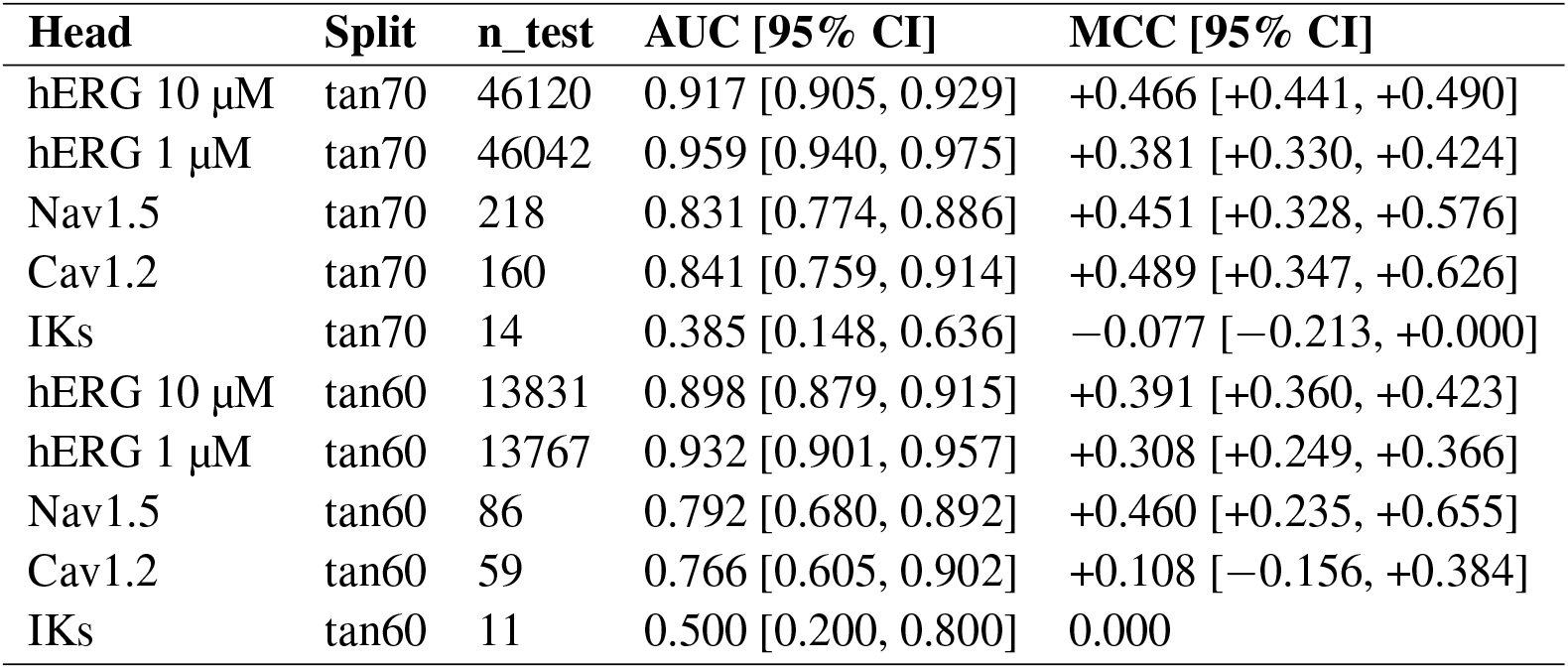
CardioSafe ensemble classification performance.

**Table 3:** CardioSafe ensemble regression performance.

| Head | Split | n_test | Pearson *r* [95% CI] | Spearman rho | RMSE [95% CI] |
| --- | --- | --- | --- | --- | --- |
| hERG pChEMBL | tan70 | 2263 | 0.553 [0.513, 0.594] | 0.553 | 0.721 [0.679, 0.769] |
| Nav1.5 pChEMBL | tan70 | 184 | 0.558 [0.429, 0.667] | 0.592 | 0.572 [0.475, 0.683] |
| Cav1.2 pChEMBL | tan70 | 124 | 0.689 [0.528, 0.811] | 0.560 | 0.815 [0.663, 0.986] |
| hERG pChEMBL | tan60 | 1205 | 0.331 [0.231, 0.499] | 0.457 | 1.001 [0.761, 1.269] |
| Nav1.5 pChEMBL | tan60 | 64 | 0.197 [-0.198, 0.704] | 0.580 | 0.838 [0.438, 1.272] |
| Cav1.2 pChEMBL | tan60 | 40 | 0.569 [0.053, 0.818] | 0.313 | 0.894 [0.713, 1.082] |

### 3.9 Named-drug case study

An 11-drug panel spanning withdrawn drugs, safe multichannel blockers, and black-box QT drugs was evaluated on tan70.

All five true hERG blockers (terfenadine, cisapride, astemizole, sertindole, domperidone) are correctly called as blockers. Verapamil is correctly flagged as a hERG blocker (0.889) with moderate Cav1.2 CO (0.584), consistent with its known dual-channel pharmacology. Ranolazine illustrates the importance of applicability domain: on tan70 (where ranolazine is out-of-train, max Tanimoto 0.588) it scores 0.123 at hERG 10 µM, while on tan60 (where it is in-train) it scores 0.719, purely because of training-set composition. The literature pIC50 for hERG–ranolazine is approximately 5.4, placing it near the 10 µM threshold; both predictions bracket the truth, but the result underscores that model output is sensitive to applicability domain.

## 4 Discussion

We presented CardioSafe, a three-branch multitask neural network with cross-attention fusion that jointly predicts blocker status and potency for hERG, Nav1.5, Cav1.2, and IKs. Four aspects of this work merit critical discussion.

### Predicted transcriptomic features: a well-characterized negative result

The integration of predicted L1000 gene expression signatures was motivated by prior evidence that transcriptional profiles can identify hERG inhibitors among structurally dissimilar compounds [34] and improve adverse drug reaction prediction [33]. Our comprehensive ablation (Tables 4 and 4b) establishes that this premise does not extend to cardiac ion channel prediction at the current data scale: neither the predicted L1000 branch nor the pretrained ChemBERTa embedding measurably improves headline classification AUC. The bio branch’s contribution is statistically indistinguishable from zero on every classification head with n ≥ 86, and on the harder tan60 split it is mildly net-negative for Cav1.2.

**Figure 4:**
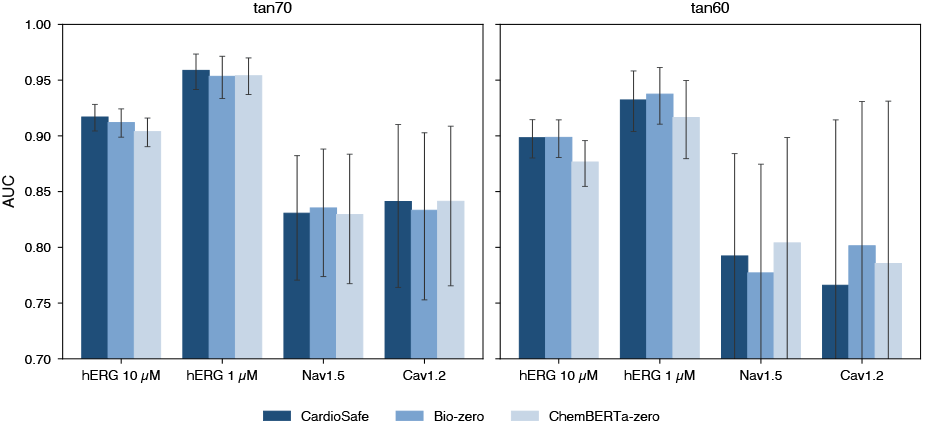
Branch ablation: ensemble AUC for the deployed CardioSafe model (all three input branches active) compared with ablations holding the L1000 biological branch to zero (bio-zero) or the ChemBERTa embedding to zero (ChemBERTa-zero), trained from scratch under the same 5 seeds and schedule. Error bars are bootstrap 95% confidence intervals (B = 1000). No ablation 95% CI falls outside the deployed model’s CI; the figure carries no asterisks, consistent with the headline negative result. The chemical fingerprint branch carries the discriminative signal; neither auxiliary branch measurably improves headline classification AUC.

**Table 4:** Pre-de-leak head-to-head classification performance on tan70 (full per-channel valid test fold)

| Channel | n | CardioSafe AUC | CToxPred2 AUC | CardioGenAI AUC | Δ-AUC vs CToxPred2 [95% CI] | Δ-AUC vs CardioGenAI [95% CI] |
| --- | --- | --- | --- | --- | --- | --- |
| hERG 10 μM | 46120 | 0.917 | 0.832 | 0.862 | +0.085 [+0.070, +0.100] | +0.055 [+0.040, +0.070] |
| Nav1.5 | 218 | 0.831 | 0.784 | 0.775 | +0.047 [−0.014, +0.107] | +0.057 [−0.013, +0.126] |
| Cav1.2 | 160 | 0.841 | 0.815 | 0.873 | +0.026 [−0.065, +0.111] | −0.032 [−0.105, +0.039] |

| Channel | CardioSafe MCC | CToxPred2 MCC | CardioGenAI MCC | Δ-MCC vs CToxPred2 [95% CI] | Δ-MCC vs CardioGenAI [95% CI] |
| --- | --- | --- | --- | --- | --- |
| hERG 10 μM | +0.466 | +0.123 | +0.147 | +0.342 [+0.322, +0.363] | +0.320 [+0.299, +0.342] |
| Nav1.5 | +0.451 | +0.384 | +0.376 | +0.068 [−0.059, +0.198] | +0.082 [−0.054, +0.211] |
| Cav1.2 | +0.488 | +0.471 | +0.516 | +0.017 [−0.153, +0.181] | −0.027 [−0.189, +0.119] |

**Table 5:** Post-de-leak head-to-head classification performance on tan70 (test fold minus comparator-Train Tanimoto ≥0.99)

| Channel | n (kept) | leak removed | Δ-AUC vs CToxPred2 [95% CI] | Δ-AUC vs CardioGenAI [95% CI] |
| --- | --- | --- | --- | --- |
| hERG 10 μM | 44814 | 2.8% | +0.131 [+0.107, +0.155] | +0.115 [+0.088, +0.143] |
| Nav1.5 | 170 | 22.0% | +0.061 [−0.002, +0.134] | +0.075 [+0.003, +0.146] |
| Cav1.2 | 127 | 20.6% | +0.106 [+0.032, +0.194] | +0.022 [−0.050, +0.100] |

| Channel | Δ-MCC vs CToxPred2 [95% CI] | Δ-MCC vs CardioGenAI [95% CI] |
| --- | --- | --- |
| hERG 10 μM | +0.327 [+0.298, +0.353] | +0.314 [+0.284, +0.342] |
| Nav1.5 | +0.091 [−0.053, +0.240] | +0.113 [−0.033, +0.254] |
| Cav1.2 | +0.146 [−0.022, +0.321] | +0.063 [−0.092, +0.221] |

**Table 6:** Branch ablation on tan70 (5-seed ensemble AUC [95% CI])

| Head | n | Deployed | Bio-zero | ChemBERTa-zero |
| --- | --- | --- | --- | --- |
| hERG 10 μM | 46120 | 0.917 [0.905, 0.929] | 0.912 [0.900, 0.924] | 0.904 [0.892, 0.916] |
| hERG 1 μM | 46042 | 0.959 [0.940, 0.975] | 0.953 [0.934, 0.973] | 0.954 [0.938, 0.971] |
| Nav1.5 | 218 | 0.831 [0.774, 0.886] | 0.835 [0.777, 0.886] | 0.829 [0.769, 0.882] |
| Cav1.2 | 160 | 0.841 [0.759, 0.914] | 0.833 [0.755, 0.905] | 0.841 [0.765, 0.913] |
| IKs | 14 | 0.385 [0.148, 0.636] | 0.462 [0.167, 0.750] | 0.385 [0.154, 0.667] |

**Table 7:** Branch ablation on tan60 (5-seed ensemble AUC [95% CI])

| Head | n | Deployed | Bio-zero | ChemBERTa-zero |
| --- | --- | --- | --- | --- |
| hERG 10 μM | 13831 | 0.898 [0.879, 0.915] | 0.899 [0.879, 0.914] | 0.877 [0.856, 0.897] |
| hERG 1 μM | 13767 | 0.932 [0.901, 0.957] | 0.937 [0.909, 0.961] | 0.916 [0.879, 0.947] |
| Nav1.5 | 86 | 0.792 [0.680, 0.892] | 0.777 [0.662, 0.881] | 0.804 [0.694, 0.899] |
| Cav1.2 | 59 | 0.766 [0.605, 0.902] | 0.801 [0.651, 0.923] | 0.785 [0.633, 0.929] |

The cross-attention layer, designed to enable per-channel specialization across the three input branches, operates at near-uniform entropy (1.098 nats vs. the maximum 1.099 nats) on every channel. No channel has learned to preferentially attend to the biological branch. The model behaves as a single-trunk chemical MLP with a small uniform-attention bonus.

We attribute this negative result to two factors. First, the LINCS L1000 panel contains no cardiac-origin cell line; the biological-branch signal is a cross-tissue perturbation signature averaged across non-cardiac cell types, and the model cannot access cardiac-tissue-specific transcriptomic context. Second, the L1000 encoder’s predicted-output graph is 29*×* denser than its training-time graph at the same co-expression threshold (47% versus 1.6% edge density), indicating that the per-gene feed-forward heads collapse predicted gene values onto a low-rank manifold, destroying the gene–gene sparsity the GCN was designed to preserve.

**Figure 5:**
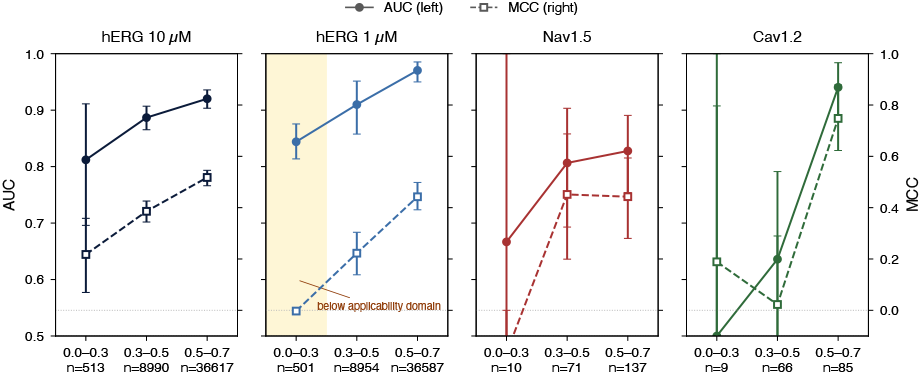
Applicability domain analysis: classification performance as a function of nearest-neighbour Tanimoto similarity to the training pool, for each of the four cardiac ion channels on the tan70 split. Each test compound is binned by its maximum Morgan-r=2 2048-bit Tanimoto to any training compound. Solid lines and filled circles show AUC (left axis); dashed lines and open squares show Matthews correlation coefficient (right axis); error bars are bootstrap 95% confidence intervals (B = 1000). Per-bin sample sizes are annotated under each tick. The hERG 1 µM head at Tanimoto < 0.3 illustrates the operational risk of relying on AUC alone: AUC remains 0.844 while MCC at the default operating point collapses to *™*0.003.

We hypothesize that cardiac-tissue-specific transcriptomic data could change this finding, although it is also possible that cross-tissue averaging provides a denoising benefit that partially offsets the tissue mismatch. The current analysis should be interpreted cautiously with respect to the expected direction of change under cardiac-specific data. Future work integrating induced pluripotent stem cell-derived cardiomyocyte (iPSC-CM) L1000 profiles would provide a direct test.

| Head | Bin | n | AUC [95% CI] | MCC [95% CI] |
| --- | --- | --- | --- | --- |
| hERG 10 μM | 0.0–0.3 | 513 | 0.812 [0.696, 0.911] | +0.218 [+0.070, +0.358] |
| hERG 10 μM | 0.3–0.5 | 8990 | 0.887 [0.866, 0.907] | +0.386 [+0.341, +0.426] |
| hERG 10 μM | 0.5–0.7 | 36617 | 0.920 [0.904, 0.935] | +0.518 [+0.489, +0.546] |
| hERG 1 μM | 0.0–0.3 | 501 | 0.844 [0.812, 0.876] | −0.003 [−0.007, +0.000] |
| hERG 1 μM | 0.3–0.5 | 8954 | 0.910 [0.862, 0.953] | +0.223 [+0.143, +0.307] |
| hERG 1 μM | 0.5–0.7 | 36587 | 0.970 [0.949, 0.986] | +0.443 [+0.385, +0.496] |
| Nav1.5 | 0.3–0.5 | 71 | 0.806 [0.695, 0.907] | +0.452 [+0.179, +0.674] |
| Nav1.5 | 0.5–0.7 | 137 | 0.828 [0.756, 0.896] | +0.443 [+0.289, +0.594] |
| Cav1.2 | 0.3–0.5 | 66 | 0.636 [0.467, 0.783] | +0.023 [−0.200, +0.271] |
| Cav1.2 | 0.5–0.7 | 85 | 0.940 [0.881, 0.985] | +0.748 [+0.624, +0.864] |

| Bucket | Count | % |
| --- | --- | --- |
| (i) Prodrug motif (hydrolyzable group) | 32 | 12% |
| (ii) Multi-target liability (blocker on another channel) | 13 | 5% |
| (iii) Label noise (>1 log-unit pChEMBL disagreement or physically impossible label) | 6 | 2% |
| (iv) Outside applicability domain (max Tanimoto < 0.30) | 8 | 3% |
| (v) True in-domain model failure | 111 | 41% |
| Borderline (max Tanimoto 0.30–0.50) | 99 | 37% |

### Evaluation methodology and the reverse-leak audit

Our Tanimoto-similarity-controlled splitting at fixed cutoffs directly guarantees that no test-set compound shares a near-neighbour with any training compound. The reverse-leak audit revealed that comparators evaluated on our test pool are contaminated: 22% of Nav1.5 and 21% of Cav1.2 test compounds appear in comparator training data, with 92% of the contamination being exact InChI-key matches and only 0.3% fuzzy near-neighbours. This issue is structural rather than procedural: all cardiac ion channel models draw from overlapping ChEMBL records, and no established protocol exists for auditing cross-publication compound overlap. Our reverse-leak analysis is intended to fill that methodological gap rather than to criticize the affected models, which followed the evaluation norms standard at the time of their publication.

### Curation policy

The pharmacology-aware curation policy that retains censored measurements and inhibition-percentage votes is validated by the strict ablation: dropping to exact-pIC50-only training costs 0.18 hERG AUC on tan70 (0.917 to 0.737). This is a load-bearing data source for binary classification, with 93% of hERG binary labels deriving from non-exact evidence. However, regression accuracy on the harder tan60 split improves under the strict policy (hERG *r*: 0.331 to 0.443), indicating the noisy measurements help classification but hurt regression. Future work could explore heteroscedastic loss functions that down-weight uncertain labels in the regression heads while preserving their contribution to classification.

### Limitations

Several limitations should be acknowledged. First, the ionization-state descriptors (pKa, logD, microstate fractions) used here are derived from a graph-based pKa predictor (MolGpKa [45]) and a Henderson–Hasselbalch independent-site model with neutral-microstate-only partitioning; conformer-aware ionization, quantum-mechanical pKa for outlier scaffolds, and explicit consideration of multiple binding-relevant microstates remain plausible refinements. Second, label noise from heterogeneous assay conditions imposes an artificial ceiling on predictive accuracy [25]. Third, the model relies on fixed fingerprint representations; graph neural networks with learned representations could be integrated as an alternative chemical branch. Fourth, the model predicts static blocker labels and pChEMBL values, which serve as inputs to the CiPA mechanistic model but are not substitutes for it. The practical deployment use case is prioritization of compounds for experimental IC50 measurement, not regulatory risk calling. Fifth, the cross-attention architecture did not learn per-channel specialization at this data scale; a simpler single-trunk multi-task model may achieve equivalent performance with fewer parameters.

| Compound | hERG pIC50 | hERG 10 μM CO | hERG 1 μM CO |
| --- | --- | --- | --- |
| Terfenadine (withdrawn, strong hERG blocker) | 6.38 ± 0.12 | 0.895 ± 0.020 | 0.763 ± 0.036 |
| Fexofenadine (safe metabolite) | 4.80 ± 0.21 | 0.683 ± 0.029 | 0.389 ± 0.056 |
| Cliff resolution (Terf – Fex) | +1.58 | +0.212 | +0.374 |

**Table 8:** CardioSafe 11-drug panel (tan70, 5-seed ensemble)

| Drug | Class | In train? | max Tan | hERG pIC50 | hERG 10 μM CO | Nav1.5 CO | Cav1.2 CO |
| --- | --- | --- | --- | --- | --- | --- | --- |
| Terfenadine | withdrawn | no | 0.891 | 6.38 | 0.895 | 0.593 | 0.627 |
| Fexofenadine | safe metabolite | no | 0.820 | 4.80 | 0.683 | 0.339 | 0.362 |
| Cisapride | withdrawn | no | 0.627 | 7.17 | 0.906 | 0.505 | 0.361 |
| Astemizole | withdrawn | yes | 1.000 | 8.35 | 0.984 | 0.813 | 0.412 |
| Sertindole | withdrawn | yes | 1.000 | 6.84 | 0.914 | 0.520 | 0.671 |
| Grepafloxacin | withdrawn | no | 0.590 | 3.90 | 0.423 | 0.095 | 0.057 |
| Verapamil | safe multichannel | yes | 1.000 | 6.62 | 0.889 | 0.506 | 0.584 |
| Amiodarone | safe multichannel | yes | 1.000 | 5.07 | 0.716 | 0.683 | 0.714 |
| Ranolazine | safe multichannel | no | 0.588 | 4.78 | 0.123 | 0.227 | 0.358 |
| Ondansetron | black-box QT | no | 0.250 | 4.43 | 0.374 | 0.181 | 0.213 |
| Domperidone | black-box QT | no | 0.885 | 6.81 | 0.915 | 0.689 | 0.455 |

## 5 Conclusions

We presented CardioSafe, a three-branch multitask neural network that jointly predicts blocker status and potency for hERG, Nav1.5, Cav1.2, and IKs. Trained on the largest publicly reported multi-channel cardiac ion channel dataset (331127 hERG, 3160 Nav1.5, 1138 Cav1.2, and 115 Iks compounds from ChEMBL 36 and hERGCentral) and evaluated under strict Tanimoto-similarity-controlled splits, the model achieved competitive AUC values across the three primary channels. Comprehensive ablation revealed that neither the predicted L1000 transcriptomic branch nor the pretrained ChemBERTa embedding measurably improves headline classification; the chemical fingerprint branch carries the model’s discriminative signal, a negative result that establishes the inadequacy of predicted cross-tissue transcriptomic context for this task and motivates future work with cardiac-specific expression data. The reverse-leak audit revealed a previously uncharacterized source of performance inflation in cross-publication bench-marking, driven by exact compound overlap between training and evaluation pools, and should become standard practice for cardiac ion channel model evaluation. A pharmacology-aware curation policy retaining censored and inhibition-percentage measurements was validated as contributing 0.18 AUC to hERG classification, demonstrating that these lower-quality evidence types carry substantial useful signal.

## 6 List of abbreviations

AD: applicability domain
APC: automated patch clamp
AUC: area under the receiver operating characteristic curve (also AUC-ROC)
AUPRC: area under the precision–recall curve
Cav1.2: voltage-gated calcium channel subtype 1.2
ChEMBL: chemical database of bioactive molecules with drug-like properties
CI: confidence interval
CiPA: Comprehensive in Vitro Proarrhythmia Assay
ECE: expected calibration error
ECFP4: extended connectivity fingerprint, diameter 4
FFN: feed-forward network
GCN: graph convolutional network
hERG: human ether-à-go-go-related gene
IC50: half-maximal inhibitory concentration
IKr: rapid component of the delayed rectifier potassium current
IKs: slow component of the delayed rectifier potassium current (KCNQ1/KCNE1)
InChI: International Chemical Identifier
iPSC-CM: induced pluripotent stem cell-derived cardiomyocyte
LLM: large language model
MAE: mean absolute error
MCC: Matthews correlation coefficient
MEA: microelectrode array
MLP: multi-layer perceptron
Nav1.5: voltage-gated sodium channel subtype 1.5
pChEMBL: *™*log10 of molar IC50/Ki/EC50/Kd as recorded in ChEMBL
pIC50: *™*log10 of molar IC50
QSAR: quantitative structure–activity relationship
RMSE: root mean square error
ROC: receiver operating characteristic
SAR: structure–activity relationship
SMILES: Simplified Molecular Input Line Entry System
TdP: Torsade de Pointes.

## 7 Declarations

### 7.1 Availability of data and materials

Curated training, validation, and test data used in this study, including the tan70 and tan60 splits and InChI-keys for all compounds, will be deposited at https://github.com/AppliedScientific/CardioSafe-benchmark under CC-BY 4.0 and MIT license. Inference will be available via a free academic API at the journal version of this manuscript; researchers seeking earlier access for benchmarking studies can contact the corresponding author. The L1000 raw signatures used to train the expression encoder are publicly available from the Gene Expression Omnibus under accessions GSE92742 (Phase I) and GSE70138 (Phase II). The ChEMBL 36 source dump (SHA-256 b25820eef0f0481ad7712bdf4bac3b45 f354e3cbacb76be1fdbf4205d6b48fb9) is available from https://www.ebi.ac.uk/chembl/.

## Additional files

All supplementary materials (Notes S1–S2, Tables S0–S9, Figure S1, the curated labels_v1 dataset, and the tan70/tan60 split indices) are available in the project repository. Specific contents:

- **Notes:** S1 (mechanism of elevated Y-randomized hERG AUC and per-stratum analysis); S2 (full literature source list and per-`pair_id` composition for the activity-cliff curation, including the 25 anchor publications and per-split filtered cliff manifests).
- **Tables:** S0 (full 20-descriptor specification, per-column non-zero fractions, summary statistics on the production cache); S1 (per-head confusion matrices); S2 and S3 (comparator panels pre-and post-de-leak, expanded versions of Tables 3 and 3b with the hERG 1 µM head and per-`pair_id` breakdown); S5 (tan60 drug panel); S6 (failure-mode representative SMILES); S7 (full applicability-domain per-bin metrics with bootstrap CIs); S8 (L1000 threshold sensitivity sweep); S9 (curation sensitivity, strict vs. full).
- **Figures:** S1 (reliability/calibration curves for hERG across applicability-domain bins).
- **Data:** the curated `labels_v1` dataset and tan70/tan60 split indices in CSV format.

## References

[1] Jamie I. Vandenberg, Matthew D. Perry, Mark J. Perrin, Stefan A. Mann, Ying Ke, and Adam P. Hill. hERG K+ channels: structure, function, and clinical significance. Physiological Reviews, 92(3):1393–1478, 2012. doi: 10.1152/physrev.00036.2011.

[2] Philip T. Sager, Gary Gintant, J. Rick Turner, Syril Pettit, and Norman Stockbridge. Rechanneling the cardiac proarrhythmia safety paradigm: a meeting report from the Cardiac Safety Research Consortium. American Heart Journal, 167(3):292–300, 2014. doi: 10.1016/j.ahj.2013.11.004.

[3] Thomas Colatsky, Bernard Fermini, Gary Gintant, Jennifer B. Pierson, Philip Sager, Yuko Sekino, David G. Strauss, and Norman Stockbridge. The Comprehensive in vitro Proarrhythmia Assay (CiPA) initiative: update on progress. Journal of Pharmacological and Toxicological Methods, 81:15–20, 2016. doi: 10.1016/j.vascn.2016.06.002.

[4] Shetuan Zhang, Zhengfeng Zhou, Qiuming Gong, Jonathan C. Makielski, and Craig T. January. Mechanism of block and identification of the verapamil binding domain to HERG potassium channels. Circulation Research, 84 (9):989–998, 1999. doi: 10.1161/01.RES.84.9.989.

[5] Bernard Fermini, Jules C. Hancox, Najah Abi-Gerges, Matthew Bridgland-Taylor, Khuram W. Chaudhary, Thomas Colatsky, Krystle Correll, William Crumb, Bruce Damiano, Gul Erdemli, Gary Gintant, John Imredy, John Koerner, James Kramer, Paul Levesque, Zhihua Li, Anders Lindqvist, Carlos A. Obejero-Paz, David Rampe, Kohei Sawada, David G. Strauss, and Jamie I. Vandenberg. A new perspective in the field of cardiac safety testing through the Comprehensive in vitro Proarrhythmia Assay paradigm. Journal of Biomolecular Screening, 21(1):1–11, 2016. doi: 10.1177/1087057115594589.

[6] Zhihua Li, Sara Dutta, Jiansong Sheng, Phu N. Tran, Wendy Wu, Kelly Chang, Thembi Mdluli, David G. Strauss, and Thomas Colatsky. Improving the in silico assessment of proarrhythmia risk by combining hERG channel-drug binding kinetics and multichannel pharmacology. Circulation: Arrhythmia and Electrophysiology, 10 (2):e004628, 2017. doi: 10.1161/CIRCEP.116.004628.

[7] Sara Dutta, Kelly C. Chang, Kylie A. Beattie, Jiansong Sheng, Phu N. Tran, Wendy W. Wu, Min Wu, David G. Strauss, Thomas Colatsky, and Zhihua Li. Optimization of an in silico cardiac cell model for proarrhythmia risk assessment. Frontiers in Physiology, 8:616, 2017. doi: 10.3389/fphys.2017.00616.

[8] Zhihua Li, Bradley J. Ridder, Xiaomei Han, Wendy W. Wu, Jiansong Sheng, Phu N. Tran, Min Wu, Aaron Randolph, Ross H. Johnstone, Gary R. Mirams, Yuri Kuryshev, James Kramer, Caiping Wu, William J. Crumb Jr, and David G. Strauss. Assessment of an in silico mechanistic model for proarrhythmia risk prediction under the CiPA initiative. Clinical Pharmacology and Therapeutics, 105(2):466–475, 2019. doi: 10.1002/cpt.1184.

[9] Gary R. Mirams, Yi Cui, Anna Sher, Martin Fink, Jonathan Cooper, Bronagh M. Heath, Nick C. McMahon, David J. Gavaghan, and Denis Noble. Simulation of multiple ion channel block provides improved early prediction of compounds’ clinical torsadogenic risk. Cardiovascular Research, 91(1):53–61, 2011. doi: 10.1093/cvr/cvr044.

[10] James Kramer, Carlos A. Obejero-Paz, Glenn Myatt, Yuri A. Kuryshev, Andrew Bruening-Wright, Joseph S. Verducci, and Arthur M. Brown. MICE models: superior to the HERG model in predicting Torsade de Pointes. Scientific Reports, 3:2100, 2013. doi: 10.1038/srep02100.

[11] Zhihua Li, Christine Garnett, and David G. Strauss. Quantitative systems pharmacology models for a new international cardiac safety regulatory paradigm. CPT: Pharmacometrics & Systems Pharmacology, 8(6):371–379, 2019. doi: 10.1002/psp4.12423.

[12] Zhihua Li, Gary R. Mirams, Takashi Yoshinaga, Bradley J. Ridder, Xiaomei Han, Janell E. Chen, Norman L. Stockbridge, Todd A. Wisialowski, Bruce Damiano, Stefano Severi, Pierre Morissette, Peter R. Kowey, Mark Holbrook, Godfrey Smith, Randall L. Rasmusson, Mei Liu, Zhen Song, Zhilin Qu, Derek J. Leishman, Joelle Steidl-Nichols, David Rampe, Christian Tabakhoff, Simon Authier, Kylie A. Beattie, Robert Wallis, Mei-Wha Jeng, Eric Wang, Tao Yang, Wei Yu, John Imredy, Martin Tristani-Firouzi, Mariko Ono, Kohei Sawada, Mariko Yamamoto Geiger, and David G. Strauss. General principles for the validation of proarrhythmia risk prediction models. Clinical Pharmacology and Therapeutics, 107(1):102–111, 2019. doi: 10.1002/cpt.1647.

[13] Jose Vicente, Robbert Zusterzeel, Lars Johannesen, Jay Mason, Philip Sager, Vikram Patel, Philip Sager, and David G. Strauss. Mechanistic model-informed proarrhythmic risk assessment of drugs. Clinical Pharmacology and Therapeutics, 103(1):54–66, 2018. doi: 10.1002/cpt.896.

[14] Hyunho Kim and Hojung Nam. hERG-Att: self-attention-based deep neural network for predicting hERG blockers. Computational Biology and Chemistry, 87:107286, 2020. doi: 10.1016/j.compbiolchem.2020.107286.

[15] Jae Yong Ryu, Min Young Lee, Jeong Hyun Lee, Byung Ho Lee, and Kwang-Seok Oh. Deep-HIT: a deep learning framework for prediction of hERG-induced cardiotoxicity. Bioinformatics, 36(10):3049–3055, 2020. doi: 10.1093/bioinformatics/btaa075.

[16] Abdul Karim, Matthew Lee, Thomas Balle, and Abdul Sattar. CardioTox net: a robust predictor for hERG channel blockade based on deep learning meta-feature ensembles. Journal of Cheminformatics, 13:60, 2021. doi: 10.1186/s13321-021-00541-z.

[17] Hyunho Kim, Minsu Park, Ingoo Lee, and Hojung Nam. BayeshERG: a robust, reliable and interpretable deep learning model for predicting hERG channel blockers. Briefings in Bioinformatics, 23(4):bbac211, 2022. doi: 10.1093/bib/bbac211.

[18] Igor H. Sanches, Rodolpho C. Braga, Vinicius M. Alves, and Carolina H. Andrade. Enhancing hERG risk assessment with interpretable classificatory and regression models (Pred-hERG 5.0). Chemical Research in Toxicology, 37(6):910–922, 2024. doi: 10.1021/acs.chemrestox.3c00400.

[19] Tianbiao Yang, Xiaoyu Ding, Elizabeth McMichael, Frank W. Pun, Alex Aliper, Feng Ren, Alex Zhavoronkov, and Xiao Ding. Atten-hERG: a reliable and interpretable graph neural network framework for predicting hERG channel blockers. Journal of Cheminformatics, 16: 143, 2024. doi: 10.1186/s13321-024-00940-y.

[20] Weiwei Wang and Roderick MacKinnon. Cryo-EM structure of the open human ether-à-go-go-related K+ channel hERG. Cell, 169(3): 422–430, 2017. doi: 10.1016/j.cell.2017.03.048.

[21] Teresa Maria Creanza, Pietro Delre, Nicola Ancona, Giovanni Lentini, Michele Saviano, and Giuseppe Felice Mangiatordi. Structure-based prediction of hERG-related cardiotoxicity: a benchmark study. Journal of Chemical Information and Modeling, 61(9):4758–4770, 2021. doi: 10.1021/acs.jcim.1c00744.

[22] Serena Vittorio, Filippo Lunghini, Alessandro Pedretti, Giulio Vistoli, and Andrea Rosario Beccari. Ensemble of structure and ligand-based classification models for hERG liability profiling. Frontiers in Pharmacology, 14:1148670, 2023. doi: 10.3389/fphar.2023.1148670.

[23] Issar Arab, Kristof Egghe, Kris Laukens, Ke Chen, Khaled Barakat, and Wout Bittremieux. Benchmarking of small molecule feature representations for hERG, Nav1.5, and Cav1.2 cardiotoxicity prediction. Journal of Chemical Information and Modeling, 64(7):2515–2527, 2024. doi: 10.1021/acs.jcim.3c01301.

[24] Issar Arab, Kris Laukens, and Wout Bittremieux. Semisupervised learning to boost hERG, Nav1.5, and Cav1.2 cardiac ion channel toxicity prediction by mining a large unlabeled small molecule data set. Journal of Chemical Information and Modeling, 64(16):6410–6420, 2024. doi: 10.1021/acs.jcim.4c01102.

[25] Gregory W. Kyro, Matthew T. Martin, Eric D. Watt, and Victor S. Batista. CardioGenAI: a machine learning-based framework for re-engineering drugs for reduced hERG liability. Journal of Cheminformatics, 17:30, 2025. doi: 10.1186/s13321-025-00976-8.

[26] Zhenqin Wu, Bharath Ramsundar, Evan N. Feinberg, Joseph Gomes, Caleb Geniesse, Aneesh S. Pappu, Karl Leswing, and Vijay Pande. MoleculeNet: a benchmark for molecular machine learning. Chemical Science, 9(2): 513–530, 2018. doi: 10.1039/C7SC02664A.

[27] Weihua Hu, Matthias Fey, Marinka Zitnik, Yuxiao Dong, Hongyu Ren, Bowen Liu, Michele Catasta, and Jure Leskovec. Open Graph Benchmark: datasets for machine learning on graphs. In Advances in Neural Information Processing Systems, volume 33, pages 22118–22133, 2020. doi: 10.48550/arXiv.2005.00687.

[28] Kevin Yang, Kyle Swanson, Wengong Jin, Connor Coley, Philipp Eiden, Hua Gao, Angel Guzman-Perez, Timothy Hopper, Brian Kelley, Miriam Mathea, Andrew Palmer, Volker Settels, Tommi Jaakkola, Klavs Jensen, and Regina Barzilay. Analyzing learned molecular representations for property prediction. Journal of Chemical Information and Modeling, 59 (8):3370–3388, 2019. doi: 10.1021/acs.jcim.9b00237.

[29] Derek van Tilborg, Alisa Alenicheva, and Francesca Grisoni. Exposing the limitations of molecular machine learning with activity cliffs. Journal of Chemical Information and Modeling, 62(23):5938–5951, 2022. doi: 10.1021/acs.jcim.2c01073.

[30] Tiago Janela and Jürgen Bajorath. Rationalizing general limitations in assessing and comparing methods for compound potency prediction. Scientific Reports, 13:17816, 2023. doi: 10.1038/s41598-023-45086-3.

[31] Tiago Janela and Jürgen Bajorath. Uncovering and tackling fundamental limitations of compound potency predictions using machine learning models. Cell Reports Physical Science, 5(6):101988, 2024. doi: 10.1016/j.xcrp.2024.101988.

[32] Aravind Subramanian, Rajiv Narayan, Steven M. Corsello, David D. Peck, Ted E. Natoli, Xiaodong Lu, Joshua Gould, John F. Davis, Andrew A. Tubelli, Jacob K. Asiedu, David L. Lahr, Jodi E. Hirschman, Zihan Liu, Melanie Donahue, Bina Julian, Mariya Khan, David Wadden, Ian C. Smith, Daniel Lam, Arthur Liberzon, Courtney Toder, Mukta Bagul, Marek Orzechowski, Oana M. Enache, Federica Piccioni, Sarah A. Johnson, Nicholas J. Lyons, Alice H. Berger, Alykhan F. Shamji, Angela N. Brooks, Anita Vrcic, Corey Flynn, Jacqueline Rosains, David Y. Takeda, Roger Hu, Desiree Davison, Justin Lamb, Kristin Ardlie, Larson Hogstrom, Peyton Greenside, Nathanael S. Gray, Paul A. Clemons, Serena Silver, Xiaoyun Wu, Wen-Ning Zhao, Willis Read-Button, Xiaohua Wu, Stephen J. Haggarty, Lucienne V. Ronco, Jesse S. Boehm, Stuart L. Schreiber, John G. Doench, Joshua A. Bittker, David E. Root, Bang Wong, and Todd R. Golub. A next generation Connectivity Map: L1000 platform and the first 1,000,000 profiles. Cell, 171(6):1437–1452, 2017. doi: 10.1016/j.cell.2017.10.049.

[33] Zichen Wang, Neil R. Clark, and Avi Ma’ayan. Drug-induced adverse events prediction with the LINCS L1000 data. Bioinformatics, 32(15): 2338–2345, 2016. doi: 10.1093/bioinformatics/btw168.

[34] Joseph J. Babcock, Fang Du, Kun Xu, Sarah J. Wheelan, and Min Li. Integrated analysis of drug-induced gene expression profiles predicts novel hERG inhibitors. PLoS One, 8(7):e69513, 2013. doi: 10.1371/journal.pone.0069513.

[35] Thai-Hoang Pham, Yuanyuan Qiu, Jingyu Zeng, Lei Xie, and Ping Zhang. A deep learning framework for high-throughput mechanism-driven phenotype compound screening and its application to COVID-19 drug repurposing. Nature Machine Intelligence, 3(3):247–257, 2021. doi: 10.1038/s42256-020-00285-9.

[36] You Wu, Qiao Liu, Yue Qiu, and Lei Xie. Deep learning prediction of chemical-induced dose-dependent and context-specific multiplex phenotype responses and its application to personalized Alzheimer’s disease drug repurposing. PLoS Computational Biology, 18(8):e1010367, 2022. doi: 10.1371/journal.pcbi.1010367.

[37] Xiaoting Tong, Naiqian Qu, Xiaohui Kong, Shuangjia Ni, Jingyu Zhou, Kun Wang, Liang Zhang, Yuyao Wen, Jiandong Shi, Songtao Zhang, Xutong Li, Yu Cong Xie, and Mingyue Zheng. TranSiGen: deep representation learning of chemical-induced transcriptional profile for phenotype-based drug discovery. Nature Communications, 15:5378, 2024. doi: 10.1038/s41467-024-49620-3.

[38] Laura-Jayne Gardiner, Anna Paola Carrieri, James Wilshaw, Stephen Checkley, Edward O. Pyzer-Knapp, and Ritesh Krishna. Using human in vitro transcriptome analysis to build trustworthy machine learning models for prediction of animal drug toxicity. Scientific Reports, 10:9522, 2020. doi: 10.1038/s41598-020-66481-0.

[39] Yoshinobu Igarashi, Noriyuki Nakatsu, Tomoya Yamashita, Atsushi Ono, Yasuo Ohno, Tetsuro Urushidani, and Hiroshi Yamada. Open TGGATEs: a large-scale toxicogenomics database. Nucleic Acids Research, 43(D1):D921–D927, 2015. doi: 10.1093/nar/gku955.

[40] Chenxi Bai, Lan Wu, Renxiao Li, Yi Cao, Song He, and Xiaochen Bo. Machine learning-enabled drug-induced toxicity prediction. Advanced Science, 2025. doi: 10.1002/advs.202413405. Verify volume/article number at proof stage.

[41] Jonathan Alvarsson, Staffan Arvidsson McShane, Ulf Norinder, and Ola Spjuth. Predicting with confidence: using conformal prediction in drug discovery. Journal of Pharmaceutical Sciences, 110(1):42–49, 2021. doi: 10.1016/j.xphs.2020.09.055.

[42] Nicolas Bosc, Francis Atkinson, Eloy Felix, Anna Gaulton, Anne Hersey, and Andrew R. Leach. Large scale comparison of QSAR and conformal prediction methods and their applications in drug discovery. Journal of Cheminformatics, 11:4, 2019. doi: 10.1186/s13321-018-0325-4.

[43] Fang Du, Haibo Yu, Beiyan Zou, Joseph Babcock, Shunyou Long, and Min Li. hERGCentral: a large database to store, retrieve, and analyze compound-human Ether-à-go-go related gene channel interactions to facilitate cardiotoxicity assessment in drug development. Assay and Drug Development Technologies, 9(6):580–588, 2011. doi: 10.1089/adt.2011.0425.

[44] Greg Landrum. RDKit: open-source cheminformatics software. https://www.rdkit.org, 2025. Accessed 28 Apr 2026.

[45] Xiaolin Pan, Hao Wang, Cuiyu Li, John Z. H. Zhang, and Changge Ji. MolGpKa: a web server for small molecule pKa prediction using a graph-convolutional neural network. Journal of Chemical Information and Modeling, 61 (7):3159–3165, 2021. doi: 10.1021/acs.jcim.1c00075.

[46] Walid Ahmad, Elana Simon, Seyone Chithrananda, Gabriel Grand, and Bharath Ramsundar. ChemBERTa-2: towards chemical foundation models. arXiv preprint arXiv:2209.01712, 2022. doi: 10.48550/arXiv.2209.01712.

[47] Joanne Bowes, Andrew J. Brown, Jacques Hamon, Wolfgang Jarolimek, Arun Sridhar, Gareth Waldron, and Steven Whitebread. Reducing safety-related drug attrition: the use of in vitro pharmacological profiling. Nature Reviews Drug Discovery, 11(12):909–922, 2012. doi: 10.1038/nrd3845.

[48] Adam Paszke, Sam Gross, Francisco Massa, Adam Lerer, James Bradbury, Gregory Chanan, Trevor Killeen, Zeming Lin, Natalia Gimelshein, Luca Antiga, Alban Desmaison, Andreas Köpf, Edward Yang, Zach DeVito, Martin Raison, Alykhan Tejani, Sasank Chilamkurthy, Benoit Steiner, Lu Fang, Junjie Bai, and Soumith Chintala. PyTorch: an imperative style, high-performance deep learning library. In Advances in Neural Information Processing Systems, volume 32, pages 8024–8035, 2019. doi: 10.48550/arXiv.1912.01703.

[49] Fabian Pedregosa, Gaël Varoquaux, Alexandre Gramfort, Vincent Michel, Bertrand Thirion, Olivier Grisel, Mathieu Blondel, Peter Prettenhofer, Ron Weiss, Vincent Dubourg, Jake Vanderplas, Alexandre Passos, David Cournapeau, Matthieu Brucher, Matthieu Perrot, and Édouard Duchesnay. Scikit-learn: machine learning in Python. Journal of Machine Learning Research, 12:2825–2830, 2011. URL https://jmlr.org/papers/v12/pedregosa11a.html.

